# LT-FGRS: a unifying R-package for the estimation of family-based genetic liabilities at population-scale

**DOI:** 10.64898/2026.06.15.731517

**Authors:** Emil M. Pedersen, Jette Steinbach, Mathias Valstad, Henrik Ohlsson, Lucas A. Rasmussen, Espen M. Eilertsen, Kenneth S. Kendler, Bjarni J. Vilhjálmsson, Andrew J. Schork, Morten D. Krebs

## Abstract

Estimates of per-individual genetic liability from large-scale family data are routinely used in biomedical research to describe genetic etiology of traits and disorders, boost the power of gene-mapping studies, and improve risk predictions. Here we present LT-FGRS, an R package for handling population-scale pedigrees and implementing multiple state-of-the-field methods for estimating genetic liability from such data. Benchmarking in population-scale Nordic registry data demonstrates that LT-FGRS reproduces estimates from existing implementations at manageable computational cost. LT-FGRS unifies previous parallel implementations into a single framework, lowering barriers for methodological comparison and applied use.

**Availability and Implementation:** LT-FGRS is available as an R-package on CRAN. (https://CRAN.R-project.org/package=LTFGRS)

**Supplementary information:** https://emilmip.github.io/LTFGRS

## 1 Introduction

Over recent years there has been increasing interest in human biomedical research to use phenotype information on genetic relatives to estimate individual-level genetic liability to disease^1–9^. These estimates find three important use-cases: 1) They can increase power of genome-wide association studies (GWAS)^6,8,10^, 2) they can be used in prediction and risk stratification models in place of or alongside of polygenic scores (PGS).^5–7^, and 3) they offer a per individual estimate of genetic liability that can be used as an instrument in genetic epidemiological studies^2,3,6^. Due to their similarity with PGS these are often referred to as *scores*, as in Family Genetic Risk Scores (FGRS).

Estimating individual level genetic liability from pedigree data has conceptual roots in classic quantitative genetic theory^11^ and a long applied history in animal breeding^12^. The use case of modern biomedical genetics present some field-specific challenges: a focus disease (binary) outcomes^10,13^, right-censored family records^6,8^, less redundancy with molecular genetic predictors^7^, more integration with gene-mapping^6,8^. Right-censored family records present a particular challenge, as existing methods differ in how they handle partially observed relatives: some ignore censoring entirely, implicitly treating censored relatives as lifetime unaffected, while others model it explicitly using either an age-dependent liability threshold or a mixture distribution (Supplementary Table S1). Risk prediction applications present a further challenge: family phenotype information must be restricted to what was observable at the time of risk assessment, unlike GWAS where all available family data can be incorporated. Together these challenges have necessitated tailored approaches^2,5,6,8,9,13^. These tailored methods have been developed in parallel, each requiring different data structures with few helper functions for structuring input, complicating their use, integration, and comparison.

LT-FGRS is an R package that provides functions and implementation vignettes for organizing and manipulating population scale family data and implementing the multiple genetic liability estimation methods developed for human biomedical research. Below, we present an overview of the existing tools, describe our software and its functionalities, and compare the various approaches in real genealogy data. A detailed description of the specific features of the existing methods is provided in the Supplementary Table S1. Briefly, the existing methods differ in: (1) the types of relatives they include; (2) if and how they account for right-censoring; (3) how they handle for family-size and relatedness among a proband’s relatives, (4) how the underlying liability is modeled, and (5) how the liability model is fit.

## 2 Methods

The **LT-FGRS** software is available as an R-package on CRAN (https://CRAN.R-project.org/package=LTFGRS). Step-wise tutorials including simulated data are available online (https://emilmip.github.io/LTFGRS/articles). Here, we briefly describe the core features of the package, which provide per-individual estimates of genetic liability from phenotypic data on family members. The estimation consists of four steps: building the genealogy, identifying informative relatives, assigning thresholds, and estimating liabilities (Figure 1).

**Figure 1.**
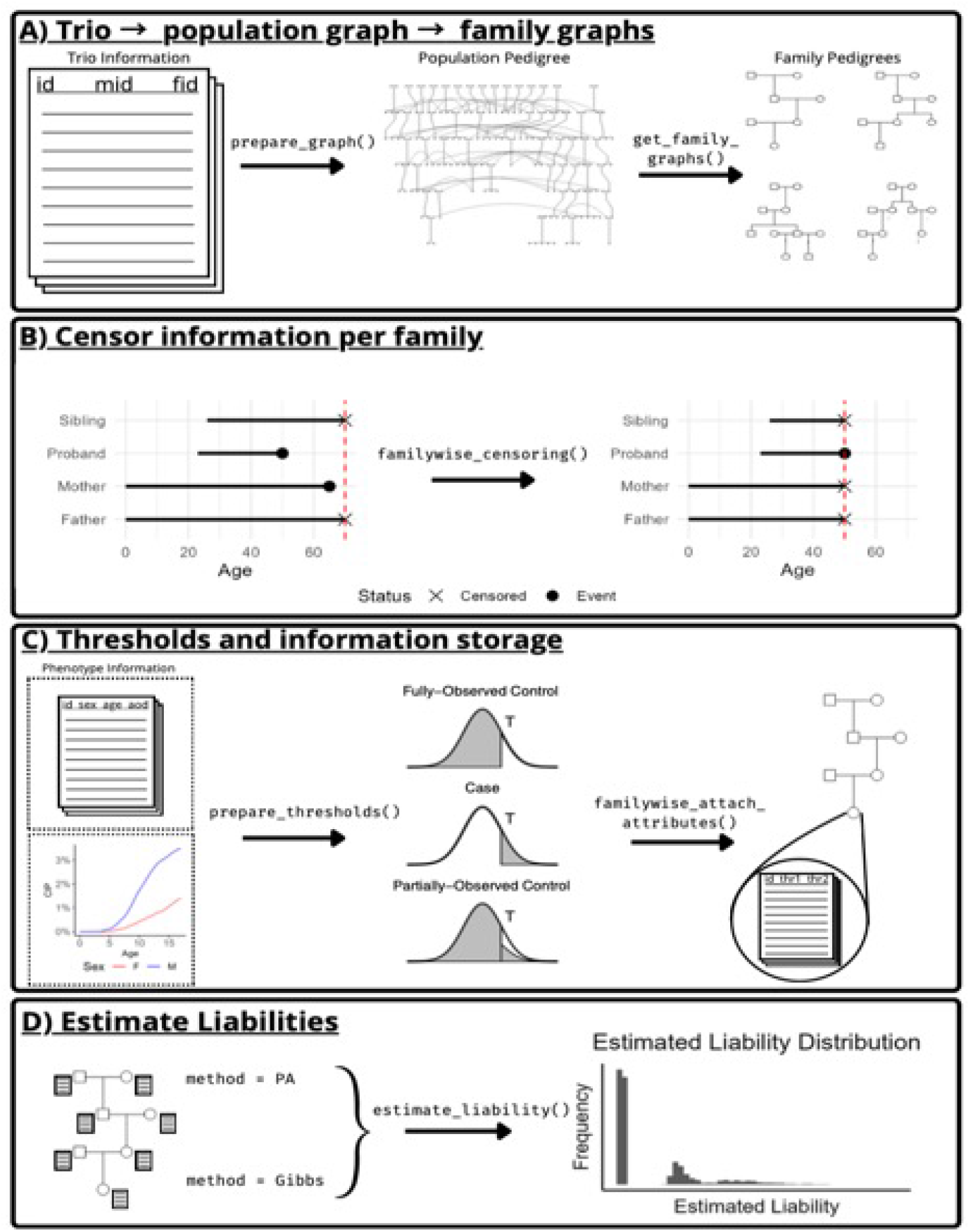
LT-FGRS workflow. The workflow of the estimating liability threshold family genetic risk scores (LT-FGRS) consist of four key steps: **A**) constructing family graphs from population level trio data. **B**) (Optional) censoring of events on a user-specified familywise basis **C**) Calculation of individualized thresholds based on population-based cumulative incidence proportions (CIP) **D**) Estimation of genetic liability scores.

### 2.1 From trios to genealogy

Family data is typically recorded as a trio record, a person identifier linked to mother and father person identifiers, which can be linked together to form pedigree tables. For population studies spanning multiple generations, this can include large, complex, highly irregular pedigrees including many relatives per individual (E.g., on average, ~22 in Denmark^6^, ~40 in Sweden^1^ and ~34 in Norway (Table S2)). LT-FGRS uses an efficient algorithm^14^ within the *prepare_graph()* function (Figure 1A) to convert trio formatted data into a *graph* object from the *igraph* software^15^. From this graph object, *get_family_graph()* can be used to construct a neighborhood graph, centred on a proband, that includes all available genetic relatives^14^.

### 2.2 Familywise censoring

For prospective risk prediction applications, the *familywise_censoring()* function (Figure 1B) censors all events within a family that happened after a specified date. The censoring date may be defined differently depending on the application, but common considerations could be based on the date of birth, age at risk assesment, age at diagnosis or date of censoring of the proband (as illustrated in Figure 1B). Such corrections have not been incorporated in other family-based liability estimation software. (For details, see https://emilmip.github.io/LTFGRS/articles/LTFGRS_workflow_prediction.html)

### 2.3 Accounting for right-censoring

LT-FGRS implements censoring corrections under the age-dependent liability threshold or the mixture model using the *prepare_thresholds()* function. This function requires estimated cumulative incidence proportions (CIP) as input, which may be stratified by sex and birth year, to generate the necessary parameters to account for censoring under either the ADT or mixture model assumptions. For details see https://emilmip.github.io/LTFGRS/articles/LTFGRS_workflow_prediction.html#assign-thresholds-to-censored-families.

### 2.4 Estimating liabilities

The *estimate_liability*() function estimates the expected genetic liability of a proband conditional on the phenotypic information of their identified relatives and a chosen censoring model. The function incorporates two ways of estimating this conditional genetic liability: 1) using a Gibbs sampler as described by^8^ and 2) using an iterative analytical approach based on the Pearson-Aitken formula as described by^6^. The censoring correction may be set to 1) no correction, 2) the ADT-, or 3) *mixture*-based correction. Finally, for comparison purposes we include a simplified implementation of the widely used FGRS_Kendler_ developed by^2^ in *simplified_kendler()*. Unlike the other methods in LT-FGRS, this approach adjusts for family structure and censoring algorithmically, rather than under an explicit model.

## 3. Results

### 3.1 Population-wide scalability

In Supplementary Table S2, we report approximate computational requirements for the estimation. While requirements may become substantial when the scores are estimated for millions of individuals with many (e.g., 34) relatives each, it’s manageable on standard HPC systems and the process is highly parallelizable.

### 3.2 New implementations are consistent with the old implementations

Figure S1 shows that we obtain liability estimates consistent with previous implementations of the included methods: Restricting to first-degree-relatives and choosing the ADT model gives estimates identical to the original LT-FH++ software. Including all relatives and choosing the *ADT* and *mixture* models, respectively, give estimates identical to PA-FGRS_ADT_ and PA-FGRS_mixt._.

### 3.3 Model choices have impact on liability estimates

Figure S1 also shows that some modelling choices are more impactful than others. Generally, estimates based on all relatives (LT-FGRS_ADT_, LT-FGRS_mixt._, LT-FGRS_ADT,FW_, LT-FGRS_mixt.,FW_, PAFGRS_ADT_, PAFGRS_mixt_, PA) are all correlated ≥0.94, and estimates based only on first-degree-relatives (LT-FGRS_1st,ADT_, LT-FGRS_1st,mixt._, LT-FGRS_1st,ADT,FW_, LT-FGRS_1st,mixt.,FW_, LTFH++, LTFH) are all correlated ≥0.97, while the correlation between these two classes are lower (0.82-0.86), showing the major differences are about which relatives they include. Ignoring censoring gives scores (PA) that are correlated 0.96 with the scores that model this LT-FGRS_ADT_ and LT-FGRS_mixt_. Familywise censoring seems to have slightly larger impact (e.g. 0.98 correlation between LT-FGRS_ADT_ and LT-FGRS_ADT,FW_,), than which model is used for handling right-censoring (e.g. 0.99 correlation between LT-FGRS_ADT_ and– LT-FGRS_mixt_).

## 4 Discussion and Conclusion

We present LT-FGRS, an R-package for estimating family-based genetic liabilities at population scale. By consolidating multiple existing methods into a single framework with standardized input formats, we aim to lower barriers for researchers seeking to leverage population-scale pedigrees and facilitate methodological comparisons.

The growing availability of nationwide health registers and large biobanks has created new opportunities for family-based genetic epidemiology. While polygenic scores have received considerable attention, family-based genetic risk scores offer a complementary approach that captures genetic liability through observed phenotypes in relatives. These two approaches have been shown to have low correlations despite similar predictive accuracies, suggesting they index partially distinct aspects of genetic risk ^5,7^. LT-FGRS provides efficient tools to compute such estimates alongside molecular genetic data.

A key contribution is the implementation of familywise censoring, ensuring that liability estimates reflect only information available at a specified index date. This is critical for valid prospective prediction, as failing to censor future family events leads to information leakage and inflated accuracy estimates.

Our results demonstrate that LT-FGRS reproduces estimates from original implementations, while offering improved scalability. The choice of which relatives to include is the primary driver of differences between methods, with estimates based on all relatives versus only first-degree relatives showing correlations of 0.82–0.86. By contrast, modeling choices around right-censoring have more modest effects.

Several limitations warrant consideration. The liability threshold models assume polygenic, normally distributed genetic architecture, which may not hold universally. The accuracy of estimates depends on phenotyping quality, which may vary by generation. Finally, interpreting family-based liabilities as purely genetic requires assuming minimal contribution from shared family environments—an assumption that may be violated for some phenotypes.

Future development could extend LT-FGRS through integration with genotype data for joint PGS-family history modeling^13^, methods for multivariate outcomes, and handling of complex family structures such as assortative mating. In conclusion, LT-FGRS provides a comprehensive framework that we hope will facilitate both methodological research and applied genetic epidemiology.

## Acknowledgements

We thank the iPSYCH consortium for providing access to the iPSYCH data. Some of the computing for this project was performed on the GenomeDK cluster. We would like to thank GenomeDK and Aarhus University for providing computational resources and support that contributed to these research results.

## Data availability

iPSYCH was approved by the Danish Scientific Ethics Committee, the Danish Health Data Authority, the Danish Data Protection Agency, Statistics Denmark, and the Danish Neonatal Screening Biobank Steering Committee. Due to the sensitive nature of the data, individual level data can only be accessed through secure servers. International researchers may obtain access through collaboration with a Danish research institution. More information about getting access can be found at https://ipsych.dk/en/about-ipsych.

