## Supplementary Figure and Tables for "LT-FGRS: a unifying R-package for the estimation of family-based genetic liabilities at population-scale"

| Method | Relatives | Model for right-censoring | Familywise censoring | Original software | LT-FGRS equivalent |
|---|---|---|---|---|---|
| **LT-FH / LT-PA** | Only 1st degree | None | - | LT-FH | *get_family_graph( degree = 1 )*<br>*prepare_thresholds( personal_thr = F )*<br>*estimate_liability( use.mixture = F )* |
| **PA** | Any | None | - | N/A | *prepare_thresholds( personal_thr = F )*<br>*estimate_liability( use.mixture = F )* |
| **PA-FGRS$_{mixt}$** | Any | mixture | - | PA-FGRS | *prepare_thresholds( personal_thr = F )*<br>*estimate_liability( use.mixture = T )* |
| **PA-FGRS$_{adt}$** | Any | ADT | - | PA-FGRS | *prepare_thresholds( personal_thr = T, lower_equal_upper = T)*<br>*estimate_liability( use.mixture = F )* |
| **LT-FH++ 1.0** | Only 1st degree | ADT | - | LTFHPlus | *get_family_graph( degree = 1 )*<br>*prepare_thresholds( personal_thr = T, lower_equal_upper = T)*<br>*estimate_liability( use.mixture = F )* |
| **FGRS$_{Kendler}$** | Any | morbid risk | - | - | kendler_simplified() |
| **LT-FGRS$_{ADT,FW}$** | Any | ADT | Yes | - | *familywise_censoring()*<br>*prepare_thresholds( personal_thr = T, lower_equal_upper = T)*<br>*estimate_liability( use.mixture=F)* |
| **LT-FGRS$_{mixt,FW}$** | Any | mixture | Yes | - | *familywise_censoring()*<br>*prepare_thresholds( personal_thr = F )*<br>*estimate_liability( use.mixture=T)* |

**Supplementary Table S1. Overview of family-based genetic liability estimation methods and their implementation in LT-FGRS.**

Each row corresponds to a method or model variant. **Relatives** indicates whether the method uses only first-degree relatives or any available relative. **Model for right-censoring** describes how partially observed relatives are handled :'None' ignores censoring, 'ADT' applies an age-dependent liability threshold, 'mixture' applies a mixture-based adjustment , and 'morbid risk' uses a weighting based on population morbid risk estimates. **Familywise censoring** indicates whether the method supports restriction of family phenotype records to a specified date, relevant for prospective risk prediction applications. **Original software** lists the previously available implementation, where applicable. **LT-FGRS equivalent** shows the function calls and parameter settings required to reproduce each method within the LT-FGRS package.

| Sample | N probands | Average family size | Memory avail. | CPU hours |
|---|---|---|---|---|
| **iPSYCH (DK)** | 141,265 | 23 | 20 GB | 3 |
| **Norwegian Population Register** | 8,966,026 | 34 | 217 GB | 19 |

**Supplementary Table S2. Computational resource usage when applying LT-FGRS to the iPSYCH and Norwegian Population Register samples.**

**N probands** indicates the number of index individuals for whom LT-FGRS was estimated. **Average family size** reflects the mean number of relatives per proband. Memory and CPU hours reflect observed usage for the runs reported here.

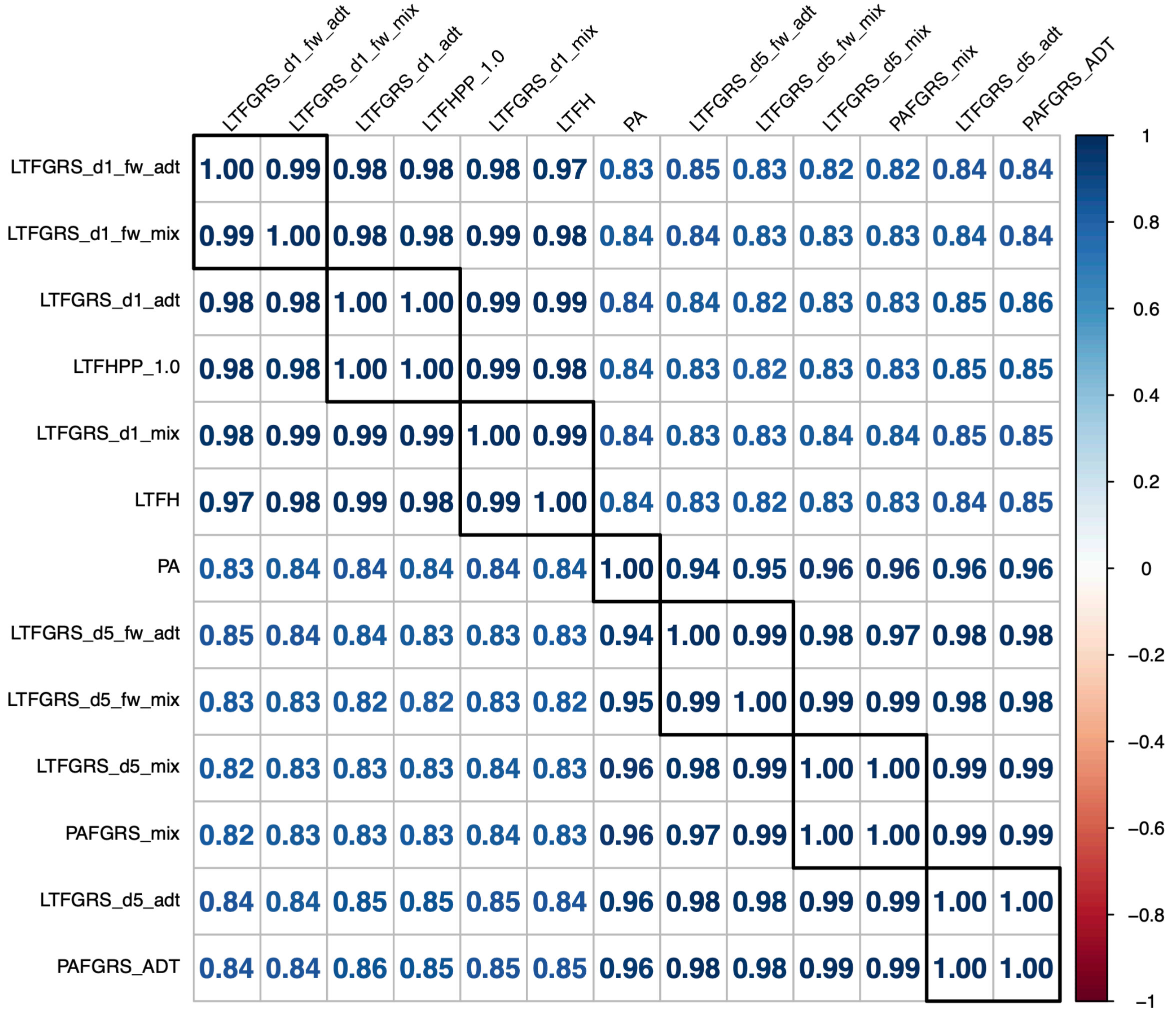


**Supplementary Figure S1 | Pairwise Pearson correlations between family genetic risk scores from different methods for major depressive disorder in the iPSYCH sample (N = 141,265).**

**LT-FGRS** are computed using 1st degree relatives only (**_d1**) or up to fifth degree relatives (**_d5**), LT-FGRS is computed without and with familywise censoring – where family members are censored at the date of diagnosis or end of follow-up of the proband (**_fw**). LT-FGRS is computed using two different right-censoring adjustments: the age-dependent-threshold (**_adt**) or the mixture (**_mix**) model. **LTFHPP_1.0** denotes the method from Pedersen et al., **LT-FH** the method from Hujoel *et al.*, **PA** the method from So *et al*., and **PA-FGRS** the method from Krebs *et al.* using the age-dependent-threshold (**_ADT**) or the mixture (**_mix**) model. Black outlines demarcate clusters of methodologically related estimators (using *hclust(k = 7)* ).
